# Cells with Treg-specific *FOXP3* demethylation but low CD25 are prevalent in autoimmunity

**DOI:** 10.1101/134692

**Authors:** Ricardo C. Ferreira, Henry Z. Simons, Whitney S. Thompson, Daniel B. Rainbow, Xin Yang, Antony J. Cutler, Joao Oliveira, Xaquin Castro Dopico, Deborah J. Smyth, Natalia Savinykh, Meghavi Mashar, Tim Vyse, David B Dunger, Helen Baxendale, Anita Chandra, Chris Wallace, John A Todd, Linda S. Wicker, Marcin L. Pekalski

**Author notes:** **Corresponding Authors:** Prof Linda S Wicker: Wellcome Trust Centre for Human Genetics, Nuffield Department of Medicine, University of Oxford, Oxford, UK. Dr Marcin L Pekalski: Wellcome Trust Centre for Human Genetics, Nuffield Department of Medicine, University of Oxford, Oxford, UK.

## Abstract

Identification of alterations in the cellular composition of the human immune system is key to understanding the autoimmune process. Recently, a subset of FOXP3^+^ cells with low CD25 expression was found to be increased in peripheral blood from systemic lupus erythematosus (SLE) patients, although its functional significance remains controversial. Here we find in comparisons with healthy donors that the frequency of FOXP3^+^ cells within CD127^low^CD25^low^ CD4^+^ T cells (here defined as CD25^low^FOXP3^+^ T cells) is increased in patients affected by autoimmune disease of varying severity, from combined immunodeficiency with active autoimmunity, SLE to type 1 diabetes. We show that CD25^low^FOXP3^+^ T cells share phenotypic features resembling conventional CD127^low^CD25^high^FOXP3^+^ Tregs, including demethylation of the Treg-specific epigenetic control region in *FOXP3* that is highly enriched in HELIOS^+^ cells, and lack of IL-2 production. As compared to conventional Tregs, more CD25^low^FOXP3^+^HELIOS^+^ T cells are in cell cycle (33.0% vs 20.7% Ki-67^+^; *P* = 1.3 x 10^-9^) and express the late-stage inhibitory receptor PD-1 (67.2% vs 35.5%; *P* = 4.0 x 10^-18^), while having reduced expression of the early-stage inhibitory receptor CTLA-4, as well as other Treg markers, such as FOXP3 and CD15s. The number of CD25^low^FOXP3^+^ T cells are highly correlated (*P* = 1.2 x 10^-19^) with the proportion of CD25^high^FOXP3^+^ T cells in cell cycle (Ki-67^+^). These findings suggest that CD25^low^FOXP3^+^ T cells represent a subset of Tregs that are derived from CD25^high^FOXP3^+^ T cells, and are a peripheral marker of recent Treg expansion in response to an autoimmune reaction in tissues.

**Highlights:** - FOXP3^+^ compartment within CD127^low^CD25^low^ T cells is expanded in autoimmune patients.

- Increased numbers of CD25^low^FOXP3^+^ T cells are a circulating marker of autoimmunity.

- CD25^low^FOXP3^+^ HELIOS^+^ T cells are fully demethylated at the *FOXP3* TSDR.

- CD25^low^FOXP3^+^ T cells could represent a terminal stage of regulatory T cells.

## 1. Introduction

FOXP3^+^ regulatory T cells (Tregs) are produced in the thymus as a specific T cell lineage following high affinity TCR engagement that results in the demethylation of the Treg-specific demethylated region (TSDR) in *FOXP3* and stable FOXP3 expression [1]. Following emigration from the thymus and activation, naïve Tregs proliferate and differentiate to memory Tregs that are actively recruited to peripheral compartments to suppress immune responses against self and maintain tissue integrity [2]. It is becoming increasingly apparent that there is considerable heterogeneity in memory Treg subsets in humans [3,4]. One major challenge for studying human Tregs is that normally only peripheral blood cells from patients are available, rather than the effector T cells and Tregs present in the inflamed tissue and associated lymph nodes. A better understanding of the composition of the Treg compartment in peripheral blood is therefore needed to investigate the potential contribution to disease mechanisms made by Tregs and to identify cellular alterations of the peripheral compartment associated with the onset of pathogenic autoimmune manifestations in the tissues.

Recently, a novel subset of FOXP3^+^ cells with low expression of CD25 was reported to be increased in peripheral blood of autoimmune systemic lupus erythematosus (SLE) patients [5–9], a finding that was later expanded to the peripheral blood of multiple sclerosis [10] and rheumatoid arthritis [11] patients. The frequency of this cell subset has been demonstrated to be associated with increased disease activity in SLE patients [5–7], suggesting that these cells may be directly pathogenic or biomarkers of flaring autoimmunity. However, the origin of these cells and their function in SLE patients and healthy individuals remain ambiguous [12,13]. In the present study, we characterise these CD127^low^CD25^low^FOXP3^+^ CD4^+^ T cells (henceforth designated as CD25^low^FOXP3^+^ cells), and demonstrate that they share phenotypic features with Tregs, including demethylation of the *FOXP3* TSDR and constitutive expression of the transcription factor HELIOS in a majority of the cells, and an inability to produce IL-2 compared to FOXP3^-^ Teffs. However, compared to conventional CD127^low^CD25^high^FOXP3^+^ Tregs, CD25^low^FOXP3^+^ cells showed increased expression of activation and proliferation markers such as PD-1 and Ki-67, and reduced expression of Treg-associated molecules, including FOXP3 and CTLA-4. We suggest that these cells represent the last stage of the natural life-cycle of TSDR-demethylated Tregs *in vivo* and that active autoimmunity increases their prevalence.

## 2. Methods

### 2.1 Subjects.

Study participants included 34 SLE patients recruited from Guy’s and St Thomas’ NHS Foundation Trust. All patients satisfied ACR SLE classification criteria and were allocated a disease activity using SLEDAI-2K at the time of sampling. SLE patients were compared to a cohort of 24 age- and sex-matched healthy donors from the Cambridge BioResource (CBR). A second cohort of 112 healthy donors from the CBR was used for the analysis of Ki-67 expression within the assessed T cell subsets.

Combined immunodeficiency patients (CID; N=7) were recruited from Cambridge University Hospitals and Papworth Hospital NHS Foundation Trusts, and compared to six age- and sex matched healthy donors from the CBR. Patients were selected on the presentation of immune infiltration in the lungs and active autoimmunity in the absence of a known genetic cause, although the clinical symptoms were consistent with those associated with recently characterised *CTLA4* germline mutations [14].

Adult long-standing T1D patients (N=15) and healthy controls (HC; N=15) were recruited from the CBR. Newly diagnosed T1D patients (ND; N=49) and unaffected siblings of other T1D probands (N=40) were collected from the JDRF Diabetes–Genes, Autoimmunity and Prevention (D-GAP) study (http://paediatrics.medschl.cam.ac.uk/research/clinical-trials/).

ND patients were characterised as having been diagnosed with T1D less than two years prior to their blood donation (with one exception of 42 months). Unaffected siblings were islet autoantibody-negative (IAA, IA2, GAD and ZnT8), and were not related to any T1D patient included in this study. All donors were of white ethnicity and all healthy controls and unaffected siblings were individuals without autoimmune disease (self-reported). Baseline characteristics for all participating subjects are summarised in **Table 1**.

### 2.2 Ethics.

All samples and information were collected with written and signed informed consent. The D-GAP study was approved by the Royal Free Hospital & Medical School research ethics committee; REC (08/H0720/25). Adult long-standing T1D patients and healthy volunteers were enrolled in the CBR. The study was approved by the local Peterborough and Fenland research ethics committee (05/Q0106/20). Informed consent was obtained from CID patients, parents, or both (R&D Ref: P01685, REC Ref: 12/WA/0148). The study conformed to the Declaration of Helsinki and all local ethical requirements.

### 2.3 PBMC sample preparation.

PBMCs were isolated by Ficoll gradient centrifugation and cryopreserved in 10% heat inactivated human AB serum, as described previously [15]. T1D patients and healthy controls were recruited contemporaneously and samples were processed and stored by the same investigators to prevent spurious findings caused by differential sample preparation.

Cryopreserved PBMCs (10x10^6^ per donor) were thawed at 37°C and resuspended in X-VIVO (Lonza) + 1% heat-inactivated, filtered human AB serum (Sigma). Cell viability following resuscitation was assessed in a subset of 40 donors using the Fixable Viability Dye eFluor 780 (eBioscience) and was found to be consistently very high (95.6%; min = 86.8%, max = 98.2%) for all samples analysed in this study.

### 2.4 Cell culture and in vitro stimulation.

To reduce the effects of experimental variation and other potential covariates, PBMC samples were processed in batches of a minimum of ten samples per day. T1D patients and healthy controls were matched as closely as possible for age (within 5 year age-bands), sex and time of sample preparation.

After thawing, PBMCs were resuspended in RPMI medium (Gibco) supplemented with 10% FBS, 2 mM L-Glutamine and 100 μg/mL Pen-Strep and cultured (10^6^ PBMCs/well) in 24 well flat-bottom cell culture plate (BD). For cytokine production assays, cells were initially rested for 30 min at 37°C and then cultured in the presence or absence of 5 ng/mL PMA, 100 ng/mL ionomycin and 0.67 μl/mL Monensin GolgiStop (BD Biosciences) for four hours at 37 °C. For a subset of 66 donors, 10^6^ cells were cultured with medium alone and 0.67 μl/mL Monensin to determine background levels of cytokine production in unstimulated cells.

### 2.5 Intracellular immunostainings.

After activation, PBMCs were harvested, and stained with Fixable Viability Dye eFluor 780 for 20 min at 4°C. Cells were then stained with fluorochrome-conjugated antibodies against surface receptors (see **Supplementary Table 1**) for one hour at 4°C. Fixation and permeabilisation was performed using FOXP3 Fix/Perm Buffer Set (BioLegend) and cells were then stained with intracellular antibodies for one hour at 4°C (see **Supplementary Table 1).** All experiments were performed in an anonymised, blinded manner without prior knowledge of disease state.

### 2.6 Flow cytometry.

*1.1* Immunostained samples were acquired using a BD Fortessa (BD Biosciences) flow cytometer with FACSDiva software (BD Biosciences) and analysed using FlowJo (Tree Star, Inc.). Dead-cell exclusion based on the Fixable Viability Dye was performed for the intracellular immunostainings.

### 2.7 Analysis of the epigenetic demethylation profile by next-generation sequencing.

Total PBMCs from seven healthy CBR donors (three males and four females) were stained with fluorophore-conjugated antibodies (see **Supplementary Table 1**) and sorted using a BD Aria Fusion flow cytometer (BD Biosciences). Methylation of the *FOXP3* TSDR was performed using a next-generation sequencing method, as described previously [16].

### 2.8 Statistical analyses.

Statistical analyses were performed using Prism software (GraphPad) and Stata (www.stata.com). Association of the assessed T-cell phenotypes with T1D, SLE and CID was calculated using two-tailed unpaired student’s t-tests. The effects of age, sex and time of collection were controlled by the experimental design used in this study and, therefore, not included as additional covariates. Given that most immune phenotypes showed moderate to strong right skew that violated the assumption of normality, the phenotypes were log transformed before statistical testing.

Comparison of the expression of the interrogated immune markers between CD127^low^CD25^low^FOXP3^+^ CD4^+^ T cells and: (i) CD25^high^FOXP3^+^, (ii) CD25^high^FOXP3^-^ and

(iii) CD25^low^FOXP3^-^ CD4^+^ T cells was performed within individuals using two-tailed paired student’s t-tests. The correlations between immune subsets were calculated using linear regression analysis.

## 3. Results

### 3.1 Frequency of CD25^low^FOXP3^+^ cells is increased in blood from patients with active autoimmunity.

To investigate the peripheral alterations in peripheral FOXP3^+^ T cell subsets, we performed a detailed immunophenotyping characterisation of cryopreserved peripheral blood mononuclear cells (PBMCs) of different cohorts of autoimmune patients (summarised in **Table 1**). Analysis of the flow cytometry profile of patients with systemic autoimmune manifestations as compared to healthy donors revealed that the frequency of FOXP3^+^ cells is highly increased in CD127^low^ cells of some patients. We found that among SLE and CID patients with increased CD127^low^ FOXP3-expressing cells there is a notable loss of CD25 expression, which results in an extremely high frequency of CD127^low^CD25^low^FOXP3^+^ cells (**Fig. 1A**). These findings suggest that the frequency of FOXP3^+^ cells in the CD127^low^CD25^low^ T cell subset (CD25^low^FOXP3^+^ cells; depicted in red in **Fig. 1B**) is increased as a result of an active autoimmune response and could be a specific marker of Treg activation. Given the lack of peripheral markers that reflect chronic immune activation, we therefore decided to focus our analysis on this population of CD25^low^FOXP3^+^ cells, and investigate their frequency in the peripheral blood of autoimmune patients.

**Fig. 1.**
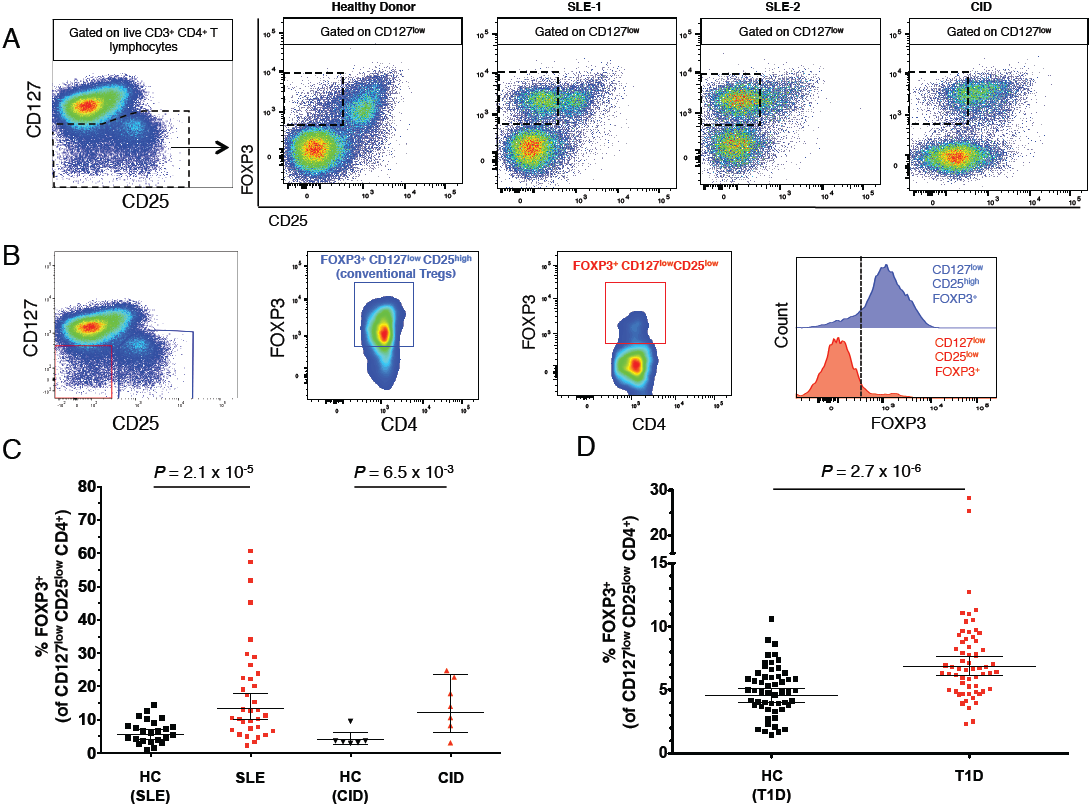
Frequency of CD25^low^FOXP3^+^ cells is increased in patients with autoimmune disease. (**A**) Patterns of CD25 and FOXP3 expression among CD127^low^ CD4^+^ T cells from healthy donors and patients with autoimmune manifestations. **(B)** Gating strategy for the delineation of the T-cell subsets characterised in this study. Distribution of FOXP3^+^ cells among: (i) CD127^low^CD25^high^ conventional Tregs (depicted in blue); and (ii) CD127^low^CD25^low^ T cells (depicted in red). The vertical dotted line represents the threshold for the gating of FOXP3^+^ cells (histograms). **(C, D)** Scatter plots depict the frequency (geometric mean +/- 95% CI) of FOXP3^+^ cells among CD127^low^CD25^low^ T cells in SLE patients (N = 32 patients vs 24 healthy donors) and combined immunodeficiency patients with active autoimmunity (N = 7 patients vs 6 healthy donors) (**C**); or in a cohort of T1D patients (N = 62; depicted by red circles) and healthy donors (N = 54; depicted by black squares) (**D**). *P* values were calculated using two-tailed unpaired t-tests. The initial CD4^+^ T cell gate (CD4 versus dead cell exclusion dye) was derived from a lymphocyte gate (defined on forward and side scatter) followed by single-cell discrimination. HC, healthy controls; T1D, type 1 diabetes patients; SLE, systemic lupus erythematosus patients; CID, combined immunodeficiency patients.

Consistent with previous findings [7,8], we confirmed that the frequency of FOXP3^+^ cells among CD127^low^CD25^low^ T cells (gating strategy **Fig. 1B**) was markedly increased in SLE patients (geometric mean (GeoM) = 13.53%) compared to age- and sex-matched healthy controls (5.51%, *P* = 2.1 x 10^-5^, N = 24, **Fig. 1C**), which likely reflects the systemic immune activation in SLE patients. In support of this hypothesis, we also detected a high frequency of CD25^low^FOXP3^+^ cells in a small cohort of seven CID patients, characterised by severe active autoimmunity compared to age- and sex-matched healthy controls (12.14% and 4.00%, respectively, *P* = 6.5 x 10^-3^; **Fig. 1C**).

We also found that the frequency of FOXP3^+^ cells among CD127^low^CD25^low^ T cells was significantly increased in T1D patients (6.84%) compared to age- and sex-matched healthy controls (4.55%; *P* = 2.7 x 10^-6^; **Fig. 1D**). This association was also observed when comparing the frequency of CD25^low^FOXP3^+^ cells within total CD4^+^ T cells (0.32% vs 0.23% in T1D patients and controls, respectively; *P* = 1.1 x 10^-3^; **Supplementary Fig. 1A**), and was not associated with duration of disease, ranging from 2 months to 23 years (*P* = 0.61). We replicated the finding of increased FOXP3^+^ cells among CD127^low^CD25^low^ T cells in an independent cohort of 15 long-standing T1D patients (10.39%) and 15 age- and sex-matched healthy controls (6.29%; *P* = 7.7 x 10^-3^; **Supplementary Fig. 1B**). Furthermore, we noted that the increased frequency of FOXP3^+^ cells was mainly restricted to the CD127^low^CD25^low^ T cell subset, as we observed only a small increased frequency of conventional CD127^low^CD25^high^FOXP3^+^ Tregs in T1D patients (5.55%) compared to healthy donors (4.82%; *P* = 8.0 x 10^-3^; **Supplementary Fig. 2**).

**Fig. 2.**
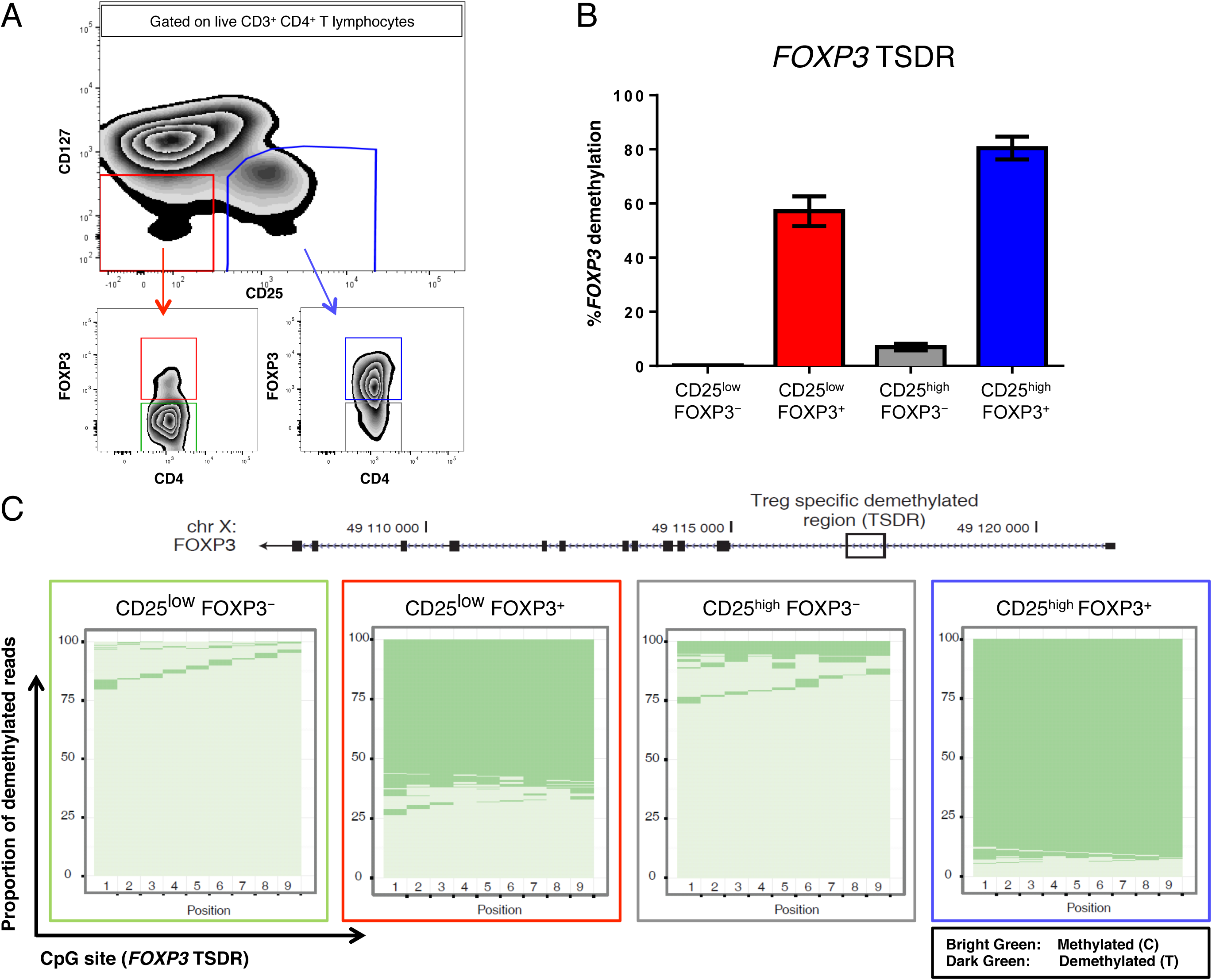
CD25^low^FOXP3^+^ cells are demethylated at the *FOXP3* Treg-specific demethylated region (TSDR). **(A)** Gating strategy for FACS sorting of four CD4^+^ T-cell subsets: (i) CD127^low^CD25^low^FOXP3^-^ (depicted in green), (ii) CD127^low^CD25^low^FOXP3^+^ (depicted in red), (iii) CD127^low^CD25^high^FOXP3^-^ (depicted in grey), and (iv) CD127^low^CD25^high^FOXP3^+^ (depicted in blue). **(B)** Frequency (mean +/- SEM) of reads demethylated at eight or nine of the nine interrogated CpG sites in the *FOXP3* TSDR. The data were obtained from sorted cells from four independent healthy donors. **(C)** Graphic depicts the proportion of demethylated reads at the nine interrogated CpG sites from the *FOXP3* TSDR in one illustrative donor. Each horizontal line represents one sequencing read, with light green representing a methylated read (C) and dark green representing a demethylated read (T). Note that the plot is representative of a male donor. For female donors, X-chromosome inactivation causes half of the reads to be methylated and a correction factor of two was applied to obtain the frequency of demethylated reads.

### 3.2 CD25^low^FOXP3^+^ cells are demethylated at the FOXP3 TSDR.

In humans FOXP3 is not exclusively expressed in Tregs, but can also be transiently up regulated in activated Teffs. However, in thymically-derived Tregs constitutive expression of *FOXP3* is known to require a demethylated TSDR [2]. To assess the TSDR methylation profile of CD25^low^FOXP3^+^ cells we sorted these cells from four healthy donors, and compared the methylation of the TSDR in CD25^low^FOXP3^+^ cells, conventional CD25^high^FOXP3^+^ Tregs and the respective FOXP3^-^ subsets (**Fig. 2A**). We found that the majority of CD25^low^FOXP3^+^ cells were demethylated at the TSDR (**Fig. 2B,C**). The epigenetic demethylation pattern in CD25^low^FOXP3^+^ cells was similar to CD25^high^FOXP3^+^ Tregs at all nine interrogated CpG sites in the *FOXP3* TSDR (mean = 57.1% and 80.5% demethylation, respectively; **Fig. 2B**); in contrast, <7% of CD25^low^FOXP3^-^ and CD25^high^FOXP3^-^ cells had demethylated TSDRs. These findings indicate the majority of CD25^low^FOXP3^+^ cells are *bona fide* Tregs, and are not Teffs transiently upregulating FOXP3 expression as a result of immune activation.

### 3.3 CD25^low^FOXP3^+^ cells express the Treg-specific HELIOS transcription factor and exhibit features of an activated phenotype.

Having established that a majority of CD25^low^FOXP3^+^ cells are stably demethylated at the *FOXP3* TSDR, we next performed a detailed phenotypic characterisation of this immune subset by flow cytometry in 24 healthy adult donors to investigate phenotypic similarities and differences between CD25^low^FOXP3^+^ and classical Tregs (CD25^high^FOXP3^+^). One distinguishing feature of these cells was the higher frequency of memory phenotype (CD45RA^-^) cells compared to their CD25^high^FOXP3^+^ counterparts (83.8% and 64.7% CD45RA^-^ cells, respectively; *P* = 2.0 x 10^-11^; **Fig. 3A**). This difference was particularly noticeable among the younger cohort (median age = 14 years) of 116 T1D patients and unaffected siblings (76.9% and 43.4%, respectively; *P* = 1.2 x 10^-58^; **Fig. 3B**), which have a higher proportion of CD45RA^+^ naïve cells amongst their CD25^high^FOXP3^+^ conventional Tregs compared to adult donors, suggesting that the majority of CD25^low^FOXP3^+^ cells have responded previously to antigen following their emigration from the thymus. Since the majority of CD25^low^FOXP3^+^ cells are CD45RA^-^, we focused further analyses on memory

**Fig. 3.**
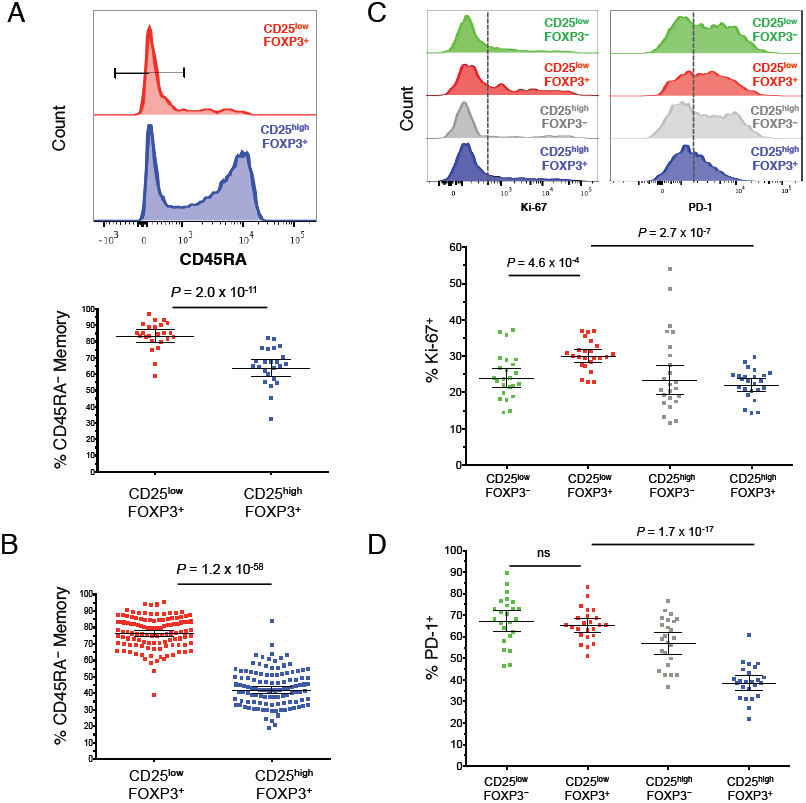
CD25^low^FOXP3^+^ T cells display an antigen-experienced phenotype. (A, B) Representative histograms and summary scatter plots depict the frequency (geometric mean +/- 95% CI) of CD45RA^-^ memory T cells amongst the CD25^low^FOXP3^-^ and CD25^low^FOXP3^+^ subsets in a population of 24 adult (median age = 42 years) healthy donors (**A**) or in a population of 116 younger (median age = 14 years) T1D patients (N = 62) and healthy donors (N = 54) (**B**). **(C, D)** Representative histograms and the frequency distribution (geometric mean +/- 95% CI) of Ki-67^+^ (**C**) and PD-1^+^ (**D**) cells in the CD45RA^-^ compartment of the four assessed immune subsets. *P* values were calculated using two-tailed paired t-tests comparing the frequency of the assessed immune subsets from the same individual. Gating strategy to delineate: (i) CD127^low^CD25^low^FOXP3^-^ (highlighted in green), (ii) CD127^low^CD25^low^FOXP3^+^ (highlighted in red), (iii) CD127^low^CD25^high^FOXP3^-^ (highlighted in grey), and (iv) CD127^low^CD25^high^FOXP3^+^ (highlighted in blue) CD4^+^ T cells is depicted in Figure 2A.

FOXP3^+^ cells.

We found that an increased frequency of CD45RA^-^ CD25^low^FOXP3^+^ cells express the proliferation marker Ki-67 compared to their CD25^high^FOXP3^+^ counterparts (29.9% and 22.0%, respectively; *P* = 2.7 x 10^-7^; **Fig. 3C**). In addition to Ki-67, CD45RA^-^ CD25^low^FOXP3^+^ cells were also characterised by a marked increased frequency of PD-1^+^ cells compared to CD25^high^FOXP3^+^ Tregs (65.1% and 38.3%, respectively; *P* = 1.7 x 10^-17^; **Fig. 3D**), and had a frequency of PD-1^+^ cells more similar to their CD25^low^FOXP3^-^ counterparts (67.2%; **Fig. 3D**).

Furthermore, we demonstrated that, similarly to CD25^high^FOXP3^+^ CD45RA^-^ memory Tregs, the majority of CD25^low^FOXP3^+^ CD45RA^-^ memory cells also express the transcription factor HELIOS, although the proportion of HELIOS^+^ cells was significantly lower (51.1%) compared to CD25^high^FOXP3^+^ CD45RA^-^ memory Tregs (77.8%; *P* = 3.0 x 10^-12^; **Fig. 4A, B)**. Consistent with the reduction in CD25 and HELIOS, CD25^low^FOXP3^+^ cells also showed a significantly lower expression of other classical Treg markers compared to CD25^high^FOXP3^+^ Tregs, such as TIGIT (65.1% vs 78.0%; *P* = 3.7 x 10^-8^), CD15s (20.7% vs 33.5%; *P* = 5.8 x 10^-12^) and most notably, CTLA-4 (60.0% vs 84.7%; *P* = 9.4 x 10^-11^; **Fig.**

**Fig. 4.**
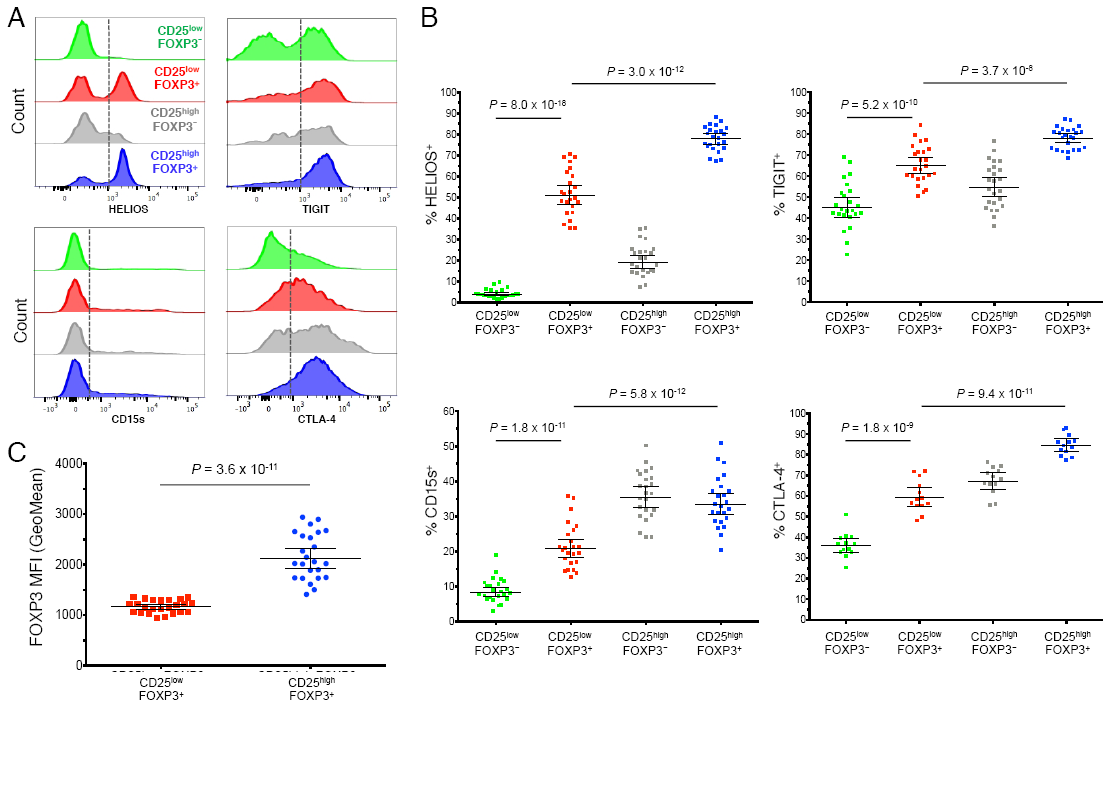
CD25^low^FOXP3^+^ cells show reduced expression of several conventional Treg markers. **(A)** Representative histograms depict the distribution of the expression of the conventional Treg markers HELIOS, TIGIT, CD15s and CTLA-4 amongst: (i) CD127^low^CD25^low^FOXP3^-^ (highlighted in green), (ii) CD127^low^CD25^low^FOXP3^+^ (highlighted in red), (iii) CD127^low^CD25^high^FOXP3^-^ (highlighted in grey), and (iv) CD127^low^CD25^high^FOXP3^+^ (highlighted in blue) memory CD4^+^ T cells. **(B)** Scatter plots depict the distribution (geometric mean +/- 95% CI) of HELIOS (n = 24), TIGIT (n = 24), CD15s (n = 24) and CTLA-4 (n = 13) in the CD45RA^-^ compartment of the four assessed immune subsets. **(C)** Expression of FOXP3 (geometric mean +/- 95% CI) was measured in the CD25^low^FOXP3^+^ (depicted by red squares) and CD25^high^FOXP3^+^ (depicted by blue circles) subsets from 24 healthy donors. *P* values were calculated using two-tailed paired t-tests comparing the assessed immunophenotypes between CD25^low^FOXP3^-^ and the other three delineated subsets from the same individual. MFI, mean fluorescence intensity.

**4A, B)**. Furthermore, we found that the expression of FOXP3 was markedly lower in CD25^low^FOXP3^+^ cells compared to CD25^high^FOXP3^+^ Tregs (MFI = 1171 and 2160 respectively, *P* = 3.6 x 10^-11^; **Fig. 4C**), suggesting that CD25^low^FOXP3^+^ cells show a decreased expression of classical Treg-associated molecules.

### 3.4 HELIOS^+^CD45RA^-^ CD25^low^FOXP3^+^ cells are demethylated at the FOXP3 TSDR to the same degree as conventional HELIOS^+^CD45RA^-^ CD25^high^ FOXP3^+^ Tregs.

To further investigate the methylation profile of FOXP3^+^ cells, we next assessed the TSDR methylation in the HELIOS^+^ and HELIOS^-^ subsets in three additional healthy donors. In agreement with their putative Treg lineage, we confirmed that the HELIOS^+^ subsets of both CD25^low^FOXP3^+^ cells and conventional CD25^high^FOXP3^+^ Tregs are virtually completely demethylated at the *FOXP3* TSDR (>95%; **Fig. 5A**). In contrast, the HELIOS^-^ subsets of CD25^low^FOXP3^+^ cells and conventional CD25^high^FOXP3^+^ Tregs showed a much lower portion of cells demethylated at the TSDR (21% and 64%, respectively). Since HELIOS expression is highly enriched in FOXP3^+^ cells demethylated at the TSDR, we examined other phenotypes within the FOXP3^+^ cells stratified by HELIOS expression. CD25^low^FOXP3^+^ cells demethylated at the TSDR as defined by HELIOS expression had a higher proportion in cell cycle as compared to CD25^high^FOXP3^+^ cells expressing HELIOS (33.0% and 20.7%, respectively; **Fig. 5B,C**). In addition, the proportion of cells expressing PD-1 and the per cell level of PD-1 were both increased on CD25^low^FOXP3^+^ demethylated at the TSDR as compared to their CD25^high^ counterparts (**Fig. 5B,C**). Expression of TIGIT, CTLA-4 and CD15s were also compared **(Supplementary Fig. 3)** with HELIOS stratification revealing a high percentage (>89%) of TIGIT^+^ cells in both populations of demethylated FOXP3^+^ cells, but a reduced number expressing CD15s and CTLA-4 in the CD25^low^FOXP3^+^ cells demethylated at the TSDR as compared to their CD25^high^ counterparts. High expression of CTLA-4 was present on CD25^high^FOXP3^+^HELIOS^-^ cells, a population with >50% of the cells having a demethylated TSDR (**Fig. 5A**). Expression of FOXP3 was found to be significantly higher within both conventional CD25^high^ Tregs (MFI > 2100) compared to CD25^low^FOXP3^+^ HELIOS^+^ T cells (MFI = 1590), despite their demethylated TSDR. Notably, the expression of FOXP3 was markedly lower in CD25^low^FOXP3^+^ HELIOS^-^ T cells (MFI = 952), which is consistent with their methylated TSDR **(Supplementary Fig. 3)**. Furthermore, analysis of CD45RA^+^ expression revealed a significantly lower frequency of CD45RA^+^ cells within total CD25^low^FOXP3^+^ HELIOS^+^ cells (4.7%) compared to their CD25^high^ counterparts (20.8%; *P* = 2.0 x 10^-12^; **Supplementary Fig. 3**). These data suggest that most of the CD45RA^+^ cells observed within CD25^low^FOXP3^+^ T cells are memory effector T cells that have re-expressed CD45RA on their surface and are characterized by being HELIOS^-^ and expressing lower levels of FOXP3. Finally, since it was possible that the expansion of CD25^low^FOXP3^+^ HELIOS^-^ cells (most of which lack a demethylated TSDR and might be activated effector cells) could have been responsible for the increase of CD25^low^FOXP3^+^ cells in autoimmune patients **(Fig. 1B,C)**, we examined the distribution of HELIOS^+^ and HELIOS^-^ cells within the CD25^low^FOXP3^+^ subset. We determined that HELIOS^+^ CD25^low^FOXP3^+^ cells were increased in SLE, CID and T1D patients as compared to their healthy control cohorts **(Supplementary Fig. 4A,B)** similar to the findings with CD25^low^FOXP3^+^ cells **(Fig. 1C,D)** and that HELIOS^+^ cells contributed significantly to all cohorts examined **(Supplementary Fig. 4C,D).**

**Fig. 5.**
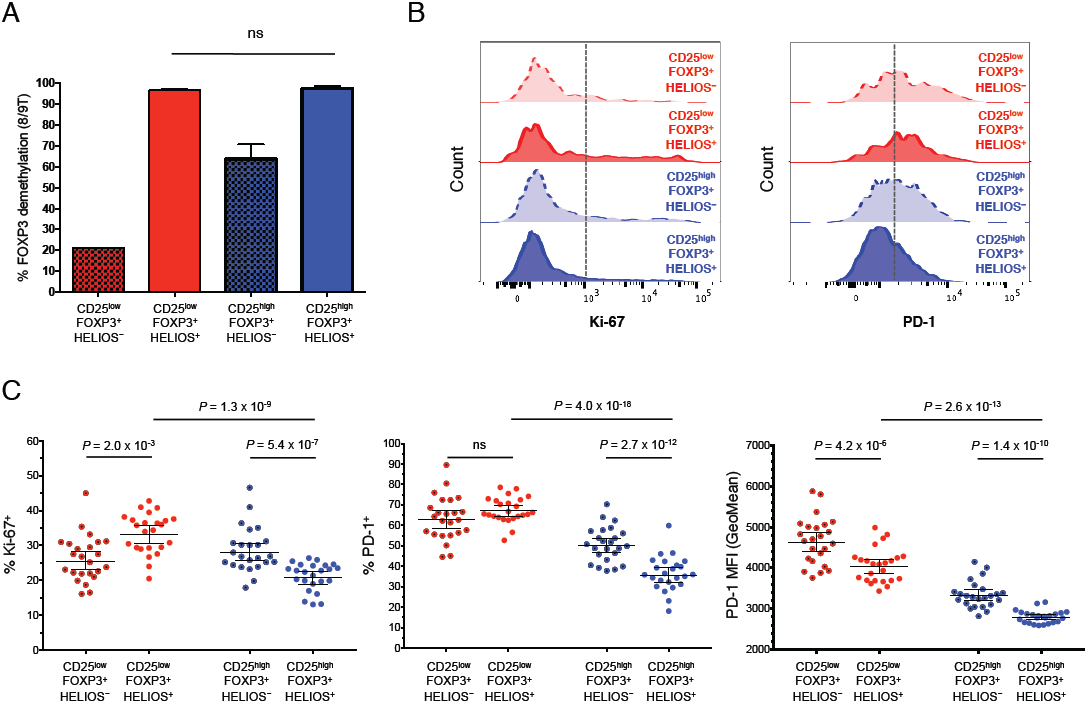
HELIOS^+^CD45RA^-^ CD25^low^FOXP3^+^ cells are demethylated at TSDR as much as conventional HELIOS^+^CD45RA^-^ CD25^high^FOXP3^+^ Tregs. Frequency (mean +/- SEM) of reads demethylated at eight or nine of the nine interrogated CpG sites in the *FOXP3* TSDR in CD45RA^-^ CD25^low^FOXP3^+^ cells and CD45RA^-^ CD25^high^FOXP3^+^ Tregs stratified by the expression of HELIOS. The data were obtained from sorted cells from three independent healthy donors.

### 3.5 Low IL-2 production from HELIOS^+^CD45RA^-^ CD25^low^FOXP3^+^ cells.

To characterise the function of HELIOS^+^CD45RA^-^ CD25^low^FOXP3^+^ cells, we assessed the production of two key cytokines, IL-2 and IFN-γ, in ten donors (five T1D patients and five healthy controls) following *in vitro* stimulation (**Fig. 6**). Consistent with their Treg-like phenotype, we found that both the HELIOS^+^CD45RA^-^ CD25^low^FOXP3^+^ and CD25^high^FOXP3^+^ subsets, which are highly demethylated at the TSDR (**Fig. 5A**), showed a low frequency of IL-2^+^ (2.0% and 1.0%, respectively) and IFN-γ^+^ cells (5.1% and 1.4%, respectively; **Fig. 6B,C**). This was in marked contrast with the HELIOS^+^CD25^low^FOXP3^-^ subset, which was found to secrete significantly higher levels of both IL-2 (21.3%; *P* = 2.8 x 10^-5^; **Fig. 6B**) and IFN-γ (27.9%; *P* = 1.1 x 10^-3^; **Fig. 6C**), compared their FOXP3^+^ counterparts (2.0% and 5.1% for IL-2 and IFN-γ, respectively). In agreement with their regulatory phenotype, we found a strong reduction of IL-2^+^ and IFN-γ^+^ cells (2.0% and 5.1%, respectively) in HELIOS^+^CD45RA^-^ CD127^low^CD25^low^FOXP3^+^ cells compared to conventional Teffs (59.8%, *P* = 5.3 x 10^-9^ and 63.5%, *P* = 2.0 x 10^-7^ for IL-2^+^ and IFN-γ^+^ cells, respectively; **Fig. 6B,C**).

**Fig. 6.**
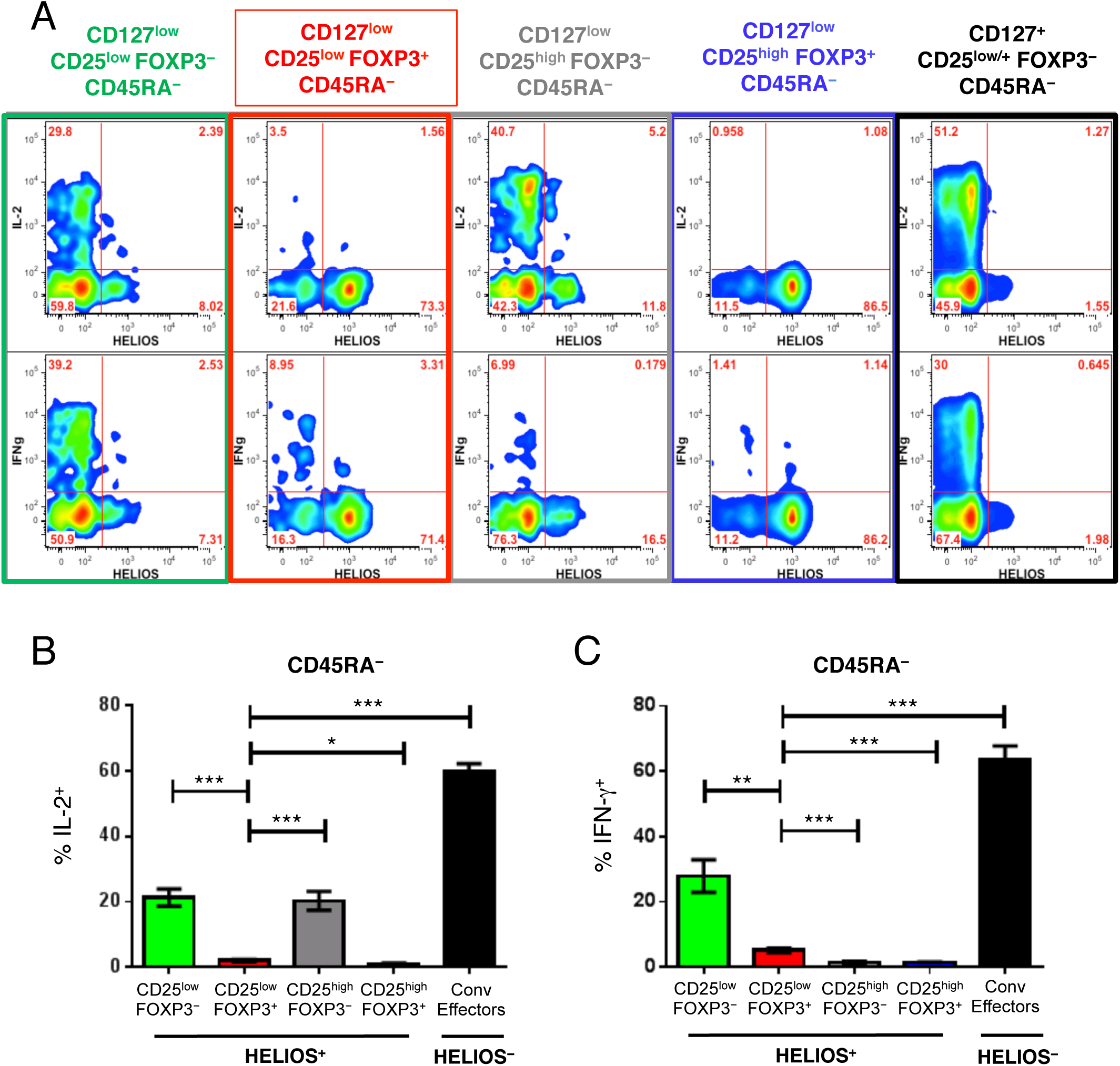
HELIOS^+^CD45RA^-^ CD25^low^FOXP3^+^ cells show impaired production of IL-2 and IFNγ. **(A)** Gating strategy to delineate the CD45RA^-^HELIOS^+^ subset of: (i) CD127^low^CD25^low^FOXP3^+^ (highlighted in red), (ii) CD127^low^CD25^high^FOXP3^+^ (highlighted in blue), and (iii) CD127^+^CD25^low/+^FOXP3^-^ HELIOS^-^ conventional (Conv) effector (highlighted in black) subsets of CD4^+^ T cells. **(B, C)** Bar graphs depict the frequency (mean +/- 95% CI) of IL-2^+^ and IFN-γ^+^ cells in the CD45RA^-^HELIOS^+^ compartment (or the CD45RA^-^HELIOS^-^ compartment in the case of the conventional effector T cells) of the five assessed immune subsets depicted in panel A. Cytokine production was assessed in one single batch of ten donors. *P* values were calculated using two-tailed paired t-tests. FACS gating plots depict data from one illustrative donor. ** *P* < 0.01, *** *P* < 0.001.

As compared to the HELIOS^+^ fraction, we found that a higher portion of HELIOS^-^ CD45RA^-^ CD25^low^FOXP3^+^ cells produced IFN-γ **(Fig. 6A, Supplementary Fig. 5A)**. These findings are consistent with a previous study, showing that HELIOS^-^FOXP3^+^ T cells produced IFN-γ, and were increased among T1D patients [17]. Although we found no evidence for differential IFN-γ production in T1D patients compared to healthy controls among HELIOS^-^ CD45RA^-^CD127^low^CD25^low^FOXP3^+^ cells, on a per cell basis (**Supplementary Fig. 5B**), the higher frequency of the CD25^low^FOXP3^+^ subset among patients resulted in a significant increase in the frequency of circulating FOXP3^+^ cells with the capability to produce IFN-γ following stimulation among total CD4^+^ T cells (*P* = 2.5 x 10^-3^; **Supplementary Fig. 5C**). These data suggest that HELIOS^-^ CD45RA^-^ CD127^low^CD25^low^FOXP3^+^ cells contributed to the increased frequency of IFN-γ^+^ cells reported among FOXP3^+^ cells from T1D patients [17].

### 3.6 CD25^low^FOXP3^+^ T cells are highly correlated with proliferating CD25^high^FOXP3^+^ Tregs.

To investigate the possible relationship between CD25^high^ and CD25^low^ FOXP3^+^HELIOS^+^ Tregs, we hypothesized that if CD25^high^ FOXP3^+^HELIOS^+^ Tregs are the precursors of the CD25^low^ FOXP3^+^HELIOS^+^ T cell subset, the numbers of CD25^high^ Tregs in cycle (Ki-67^+^) and CD25^low^ FOXP3^+^HELIOS^+^ T cells should be correlated. This correlation would be required to maintain homeostasis of Treg numbers such that as memory CD25^high^ Tregs are required to increase in peripheral compartments to respond to inflammatory conditions, a higher Treg turnover would lead to more CD25^high^ Tregs moving into the CD25^low^ compartment and ultimately to cell death. We assessed the total numbers of CD45RA^-^ Ki-67^+^ Tregs both in the cohort of 24 healthy volunteers (cohort 1) and in an independent replication cohort (cohort 2) of 112 healthy volunteers. We found that the frequency of CD45RA^-^ CD4^+^ Ki-67^+^ CD25^high^FOXP3^+^HELIOS^+^ Tregs was significantly correlated with the frequency of CD25^low^FOXP3^+^HELIOS^+^ T cells within total memory CD4^+^ T cells (r^2^ = 0.18, *P* = 3.1 x 10^-7^; **Fig. 7A**). Similarly, we observed a strong correlation between the numbers of in-cycle Ki-67^+^ Tregs and the CD25^high^FOXP3^+^HELIOS^+^ Treg compartment (r^2^ = 0.46, *P* = 1.2 x 10^-19^; **Fig. 7B**), as well as a significant correlation between both the total CD25^low^ and CD25^high^ FOXP3^+^HELIOS^+^ T cell compartments (r^2^ = 0.15, *P* = 3.9 x 10^-6^; **Fig. 7C**). These observed correlations were very consistent within both cohorts of healthy volunteers, and suggest that proliferation of conventional CD25^high^ FOXP3^+^HELIOS^+^ T cells is critical to promote the homeostatic repopulation of the CD25^high^ Treg subset, which is maintained at a steady state frequency through the progression of a proportion of CD25^high^ Tregs to the CD25^low^ FOXP3^+^HELIOS^+^ compartment.

**Fig. 7.**
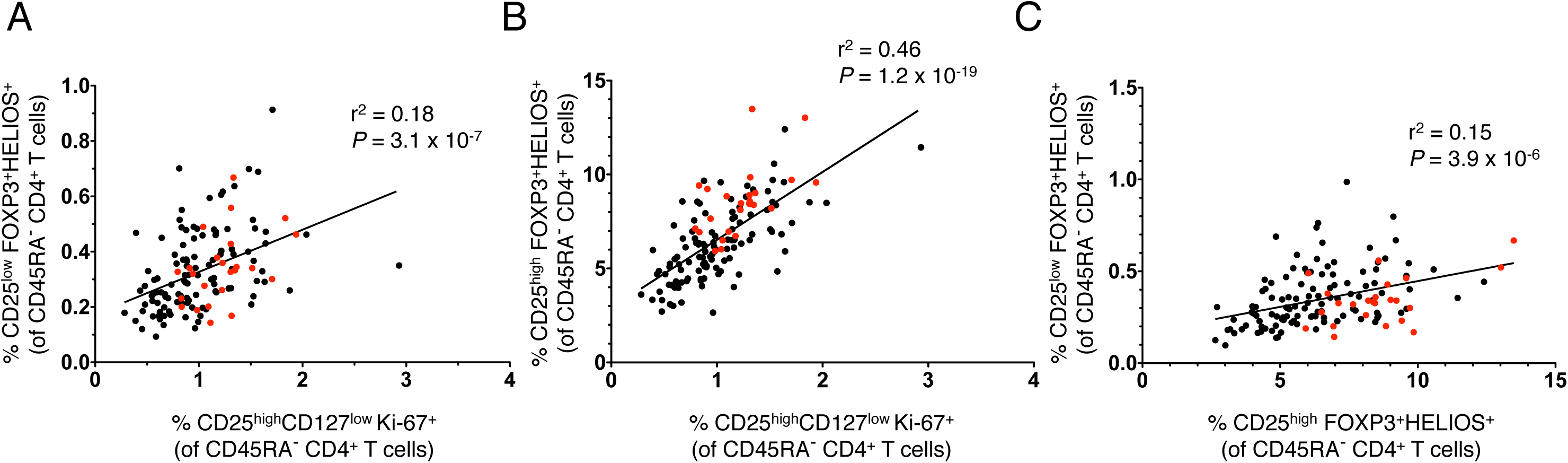
Proliferating Ki-67^+^ CD127^low^CD25^high^ Tregs correlate with the frequencies of the CD127^low^CD25^low^ and CD127^low^CD25^high^ HELIOS^+^FOXP3^+^subsets. (A, B) Data shown depict the correlation between the frequency within CD45RA^-^ CD4^+^ T cells of in-cycle (Ki-67^+^) CD4^+^CD45RA^-^ CD127^low^CD25^high^ Tregs (FOXP3^+^HELIOS^+^) and the frequency of either CD25^low^ FOXP3^+^HELIOS^+^ T cells (A) or conventional CD25^high^ FOXP3^+^HELIOS^+^ Tregs (B). (C) Data shown depict the correlation between the frequencies of circulating CD4^+^CD45RA^-^ CD25^low^ FOXP3^+^HELIOS^+^ and CD25^high^ FOXP3^+^HELIOS^+^ T cells. Frequencies of the assessed immune subsets were measured in PBMCs from healthy volunteers from two independent cohorts: cohort 1 containing 24 donors (depicted in red) and cohort 2 containing 112 donors (depicted in black). The r^2^ values represent the coefficient of determination of the linear regression in the combined cohorts, and the *P* values correspond to the F statistic testing the null hypothesis that the slope of the linear regression analysis is equal to 0.

## 4. Discussion

The identification of reliable biomarkers of disease activity has been a major challenge of autoimmune diseases, particularly in organ-specific diseases, such as T1D, where there is limited access to the inflamed tissues. In this study we characterised a subset of FOXP3^+^ CD127^low^CD25^low^ T cells, and show that it could be a peripheral biomarker of a recent autoimmune reaction in the tissues. We showed that in addition to SLE, where the increase of FOXP3^+^ CD25^low^ T cells has been observed in multiple SLE studies [5–9], the proportion of FOXP3^+^ cells in the CD127^low^CD25^low^ subset is increased in CID and T1D patients.

Although the frequency of FOXP3^+^ cells in the CD127^low^CD25^low^ subset was compared in T1D patients versus controls in one previous study with no difference observed [13], we note that the number of participants was small: 10 healthy control individuals and 16 patients. In contrast, Zoka et al [18] observed that the proportion of CD25^low^ cells among FOXP3^+^CD4^+^ T cells is higher in T1D patients than in controls, a phenotype consistent with our observations.

A major strength of this study is that we were able to use a recently developed assay [16] to precisely assess the methylation status of the *FOXP3* TSDR of CD25^low^FOXP3^+^ cells, a feature that was lacking in the previous SLE studies [7,8] or studies of this subset from healthy individuals [13]. This method provides a more quantitative assessment of the methylation pattern of the *FOXP3* locus [16] in the different immune subsets, which allowed us to demonstrate that the epigenetic profile of CD25^low^FOXP3^+^ cells was remarkably similar to conventional CD25^high^ FOXP3^+^ Tregs (57.1% and 80.5% demethylated at the TSDR, respectively). We went on to define that this epigenetic similarity was caused primarily by the TSDR methylation status of HELIOS^+^ cells: virtually all HELIOS^+^ cells were found to be demethylated in both the CD25^high^ and CD25^low^ FOXP3^+^ subsets. This finding is consistent with a previous study assessing the *FOXP3* TSDR methylation profile in different CD4^+^CD127^low^ T-cell subsets discriminated by their expression of FOXP3 and CD25 [19]. The majority of CD25^low^FOXP3^+^ cells sorted from synovial fluid mononuclear cells of juvenile idiopathic arthritis patients were shown to have a demethylated *FOXP3* TSDR, suggesting that this subset may be enriched at inflammatory sites [19]. In the current study we also found that the proportion of cells expressing TIGIT was elevated over 2-fold in both FOXP3^+^HELIOS^+^ subsets as compared to their FOXP3^+^HELIOS^-^ counterparts. Stable demethylation of the *FOXP3* TSDR occurs in the thymus upon strong T-cell receptor stimulation [1], therefore suggesting that CD25^low^ FOXP3^+^HELIOS^+^ cells are *bona-fide* thymically-derived Tregs that have lost the expression of CD25.

In further support of the hypothesis that CD25^low^FOXP3^+^ HELIOS^+^ T cells are in fact a subset of the classical FOXP3^+^ Treg subset, these cells were unable to produce IL-2 following *in vitro* activation. This finding was in contrast with the report by Yang *et al* [8] that CD25^low^FOXP3^+^ T cells from new-onset SLE patients were able to secrete IL-2.

However, we note that the IL-2 production reported in Yang *et al* was much lower compared to CD25^high^FOXP3^-^ Teffs, and immune subsets were not stratified based on the expression of CD127, CD45RA and HELIOS. It is therefore likely that the residual production of IL-2 observed by Yang *et al* in CD25^low^FOXP3^+^ T cells was due primarily to HELIOS^-^ T cells. In contrast, in our study we demonstrate that CD45RA^-^ CD25^low^CD127^low^ HELIOS^+^FOXP3^+^ cells have a profound inability to produce IL-2 as compared to CD127^+^CD25^high^ CD45RA^-^ HELIOS^-^FOXP3^-^ Teffs. We also noted the overall heterogeneity in the CD127^low^ subset in regard to IL-2 and IFN-γ secretion (**Fig. 6**). A similar proportion of CD127^low^ cells lacking both FOXP3 and HELIOS expression secrete IL-2 and IFN-γ as compared to their CD127^+^ counterparts and are likely effector T cells. In healthy individuals we observed that these putative effector cells are the largest portion of the CD45RA^-^ CD25^low^CD127^low^ gate (**Fig. 1B**), consistent with previous observations [13].

In addition to reduced levels of CD25, CD25^low^ HELIOS^+^FOXP3^+^ Tregs had lower expression of CTLA-4, CD15s and FOXP3 as compared to CD25^high^ HELIOS^+^FOXP3^+^ Tregs, suggesting that CD25^low^ Tregs could have decreased suppressive function. One limitation of our study is that we are not able to directly assess the suppressive capacity of CD25^low^ HELIOS^+^FOXP3^+^ cells, as sorting on the intracellular transcription factors precludes the use of these cells for functional assays and surrogate surface markers are not yet defined. Also, as described above, the CD45RA^-^ CD25^low^CD127^low^ gate has a high proportion of effector cells present making the results of suppression experiments using populations of cells gated as CD25^low^CD127^low^ difficult to interpret. Notably, despite the cellular heterogeneity inherent in the CD25^low^CD127^low^ subset, two studies did test the suppressive capacity of sorted CD25^low^CD127^low^ CD4^+^ T cells [7,13]. Suppression of proliferation by Teffs was mediated by CD25^low^CD127^low^ CD4^+^ T cells in both studies; however IFN-γ secretion by Teffs was not suppressed in the one study that examined this parameter [7]. The reduced suppression mediated by CD25^low^CD127^low^ CD4^+^ T cells could be due to the fact that a larger proportion of effector cells are present in this subset as compared with their CD25^+^ counterparts. Overall the observation of suppression by CD25^low^CD127^low^ CD4^+^ T cells supports the conclusion that the CD25^low^ HELIOS^+^FOXP3^+^ Tregs present in the heterogeneous CD25^low^CD127^low^ CD4^+^ T cell population are functionally suppressive. Future studies are needed to unravel the heterogeneity present in both the CD25^low^ and CD25^high^ CD127^low^CD4^+^ T cell subsets.

In contrast to the reduced expression of several molecules that are abundant in FOXP3^+^ Tregs, the frequency of cells expressing PD-1 and Ki-67 in CD25^low^ Tregs was higher than in conventional CD25^high^ Tregs, suggesting that the CD25^low^ HELIOS^+^FOXP3^+^ population may represent the consequences of CD25^high^ Tregs attempting to suppress ongoing inflammatory responses in tissues. The progression of CD25^high^ Tregs to the CD25^low^ Treg subset is supported by our observation of the strong correlation between the frequency of CD25^high^ Tregs in cell cycle (Ki-67^+^) with the number of CD25^low^ Tregs. The high proportion (15-40%) of memory FOXP3^+^ Tregs in cycle is consistent with their shorter half-lives as compared to other T cell subsets [20,21]. Thus, given the fact that Treg percentages normally remain constant in an individual through time [21], and a high proportion of the cells are replicating, Treg cell death must be a common outcome following cell division. We propose that the decreased expression of CD25 on Tregs, most likely caused by exposure to inflammatory conditions, causes less responsiveness to IL-2, reduced expression of FOXP3 and other Treg-associated molecules, and an increased probability of cell death. Despite the reduced IL-2 responsiveness in CD25^low^ Tregs, it is possible that the Tregs remain functional and that the upregulation of PD-1 could compensate for reductions in FOXP3 and CTLA-4 levels [22]. Consistent with this hypothesis, previous studies have reported that PD-1 is a critical inhibitory molecule that is upregulated on T cells after activation [23–25]. In contrast, chronic PD-1 signaling within peripheral compartments has been reported to lead to reduced STAT5 phosphorylation, decreased expression of CD25, FOXP3 and CTLA-4, and decreased

Treg suppressive function [26–28]. Additional functional studies are required to resolve these apparently contradictory mechanisms.

## 5. Conclusions

We hypothesize that the presence of a low frequency of CD25^low^ HELIOS^+^FOXP3^+^ cells in peripheral blood from healthy individuals reflects a normal physiological mechanism to maintain, genetically-regulated, Treg levels. Their increased frequency in peripheral blood from autoimmune patients, which is particularly noteworthy in patients with chronic systemic inflammation, is indicative of an inflammatory insult that drives the expansion of the Treg population, which can be transient or chronic, in an attempt to regulate an overt autoimmune Teff response. Given the paucity of reliable peripheral biomarkers of disease activity, our findings suggest that the frequency of CD25^low^ HELIOS^+^FOXP3^+^ Tregs could provide valuable information about recent or ongoing tissue inflammation and could have a clinical application for the stratification of patients with flaring autoimmunity.

## Author contributions

R.C.F., J.A.T., L.S.W. and M.L.P. designed experiments and interpreted data. R.C.F., H.Z.S., W.S.T., D.B.R., A.J.C., J.O., X.C.D., D.J.S., N.S., M.M. and M.L.P. performed experiments. X.Y. analysed the data. C.W. supervised the statistical analysis of the data. T.V., D.B.D., H.B and A.C. provided samples and clinical outcome data. R.C.F., J.A.T., L.S.W. and M.L.P. conceived the study and wrote the paper.

## Acknowledgements

This work was supported by the JDRF UK Centre for Diabetes - Genes, Autoimmunity and Prevention (D-GAP; 4-2007-1003) in collaboration with M. Peakman and T. Tree at Kings College London, the JDRF, the Wellcome Trust (WT; WT061858/091157) and the National Institute for Health Research Cambridge Biomedical Research Centre. RCF is funded by an advanced JDRF post-doctoral fellowship (3-APF-2015-88-A-N). CW is funded by the Wellcome Trust (088998).

We thank staff of the National Institute for Health Research (NIHR) Cambridge BioResource recruitment team for assistance with volunteer recruitment and K. Beer, T. Cook, S. Hall and J. Rice of the Cambridge BioResource for blood sample collection. We thank C. Guy from the Department of Paediatrics, School of Clinical Medicine, University of Cambridge for D-GAP sample recruitment. We thank M. Woodburn and T. Attwood from the Cambridge Institute for Medical Research, University of Cambridge for their contribution to sample management and N. Walker and H. Schuilenburg from the Cambridge Institute for Medical Research, University of Cambridge for data management. This research was supported by the Cambridge NIHR BRC Cell Phenotyping Hub. In particular, we wish to thank Anna Petrunkina Harrison, Simon McCullum, Christopher Bowman and Esther Perez from the Cambridge NIHR BRC Cell Phenotyping Hub for their advice and support in cell sorting. We thank Howard Martin, Fay Rodger and Ruth Littleboy for running the Illumina MiSeq in the Molecular Genetics Laboratories, Addenbrooke’s Hospital, Cambridge. We thank members of the NIHR Cambridge BioResource SAB and management committee for their support and the NIHR Cambridge Biomedical Research Centre for funding. Access to NIHR Cambridge BioResource volunteers and their data and samples is governed by the NIHR Cambridge BioResource SAB. Documents describing access arrangements and contact details are available at http://www.cambridgebioresource.org.uk/. We also thank H. Stevens, P. Clarke, G. Coleman, S. Dawson, S. Duley, M. Maisuria-Armer and T. Mistry from the Cambridge Institute for Medical Research, University of Cambridge for preparation of PBMC samples.

## Supplement

**Supplementary Table 1.** Antibodies and immunostaining panels used for flow cytometry. Detailed description of the fluorochrome-conjugated antibodies and immunostaining panels used in this study.

**Supplementary Fig. 1.**
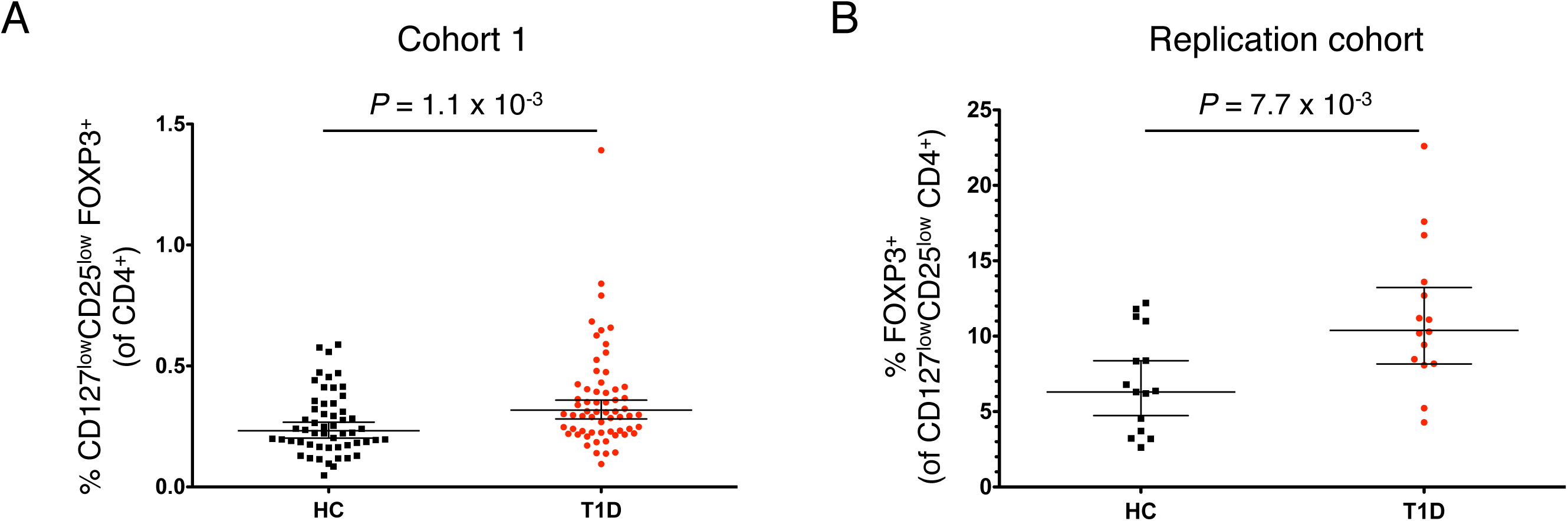
Frequency of CD127^low^CD25^low^ FOXP3^+^ T cells is increased in T1D patients. (**A**) Scatter plot depicts the total frequency (geometric mean +/- 95% CI) of CD25^low^FOXP3^+^ cells out of CD4^+^ T cells in our discovery cohort of 62 T1D patients (depicted by red circles) and 54 healthy controls (depicted by black squares) (**B**) Scatter plots depict the frequency (geometric mean +/- 95% CI) of FOXP3^+^ cells from CD127^low^CD25^low^ T cells in: (i) an independent replication cohort consisting of 15 T1D patients and 15 healthy controls. *P* values were calculated using two-tailed unpaired t-tests comparing the geometric mean of the assessed immune subsets between T1D patients and healthy controls (HC).

**Supplementary Fig. 2.**
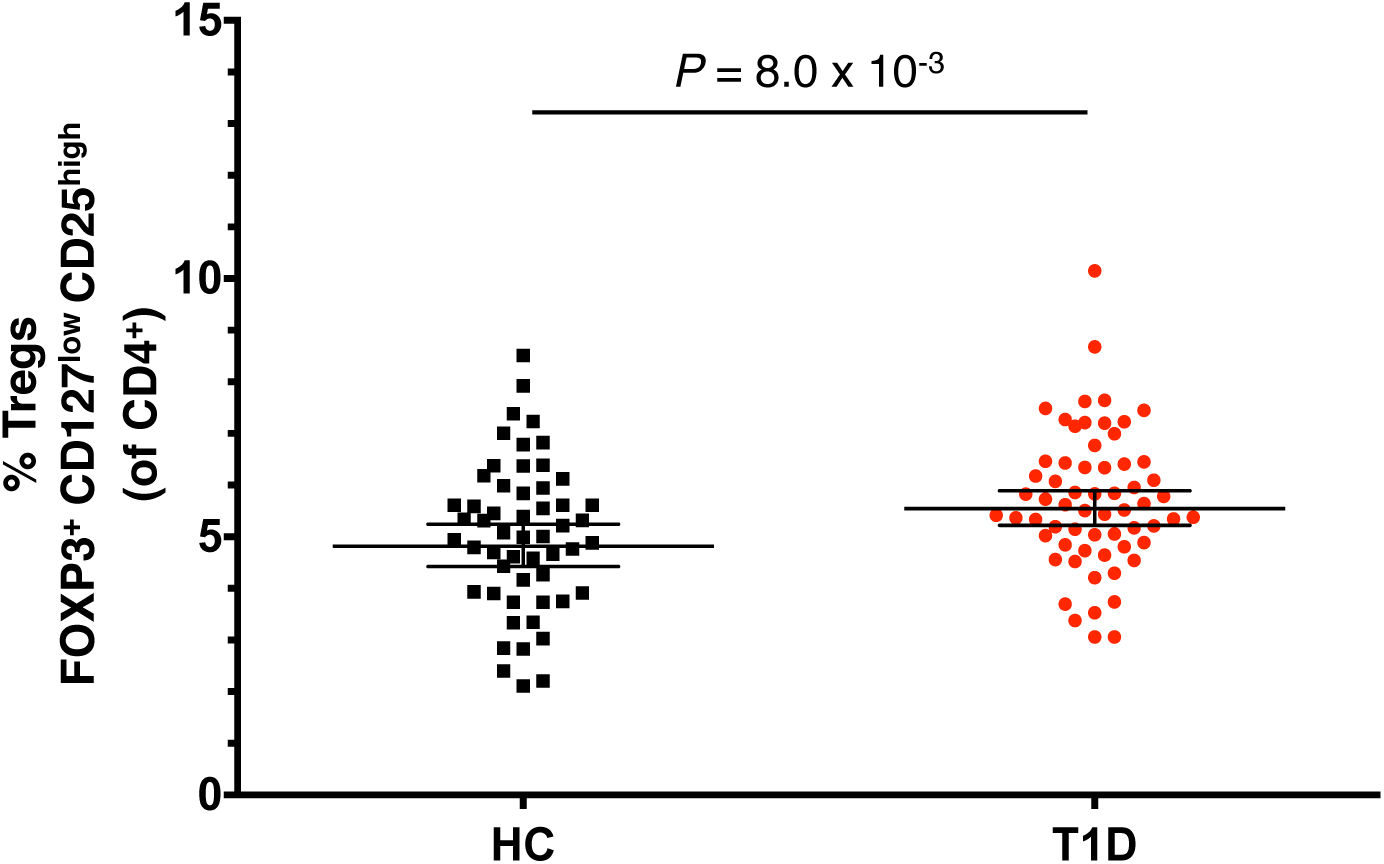
Minimal increase in the frequency of CD127^low^CD25^high^ FOXP3^+^ T cells in T1D patients. Scatter plot depicts the total frequency (geometric mean +/- 95% CI) of CD25^high^FOXP3^+^ cells (classical Tregs) out of CD4^+^ T cells in our discovery cohort of 62 T1D patients (depicted by red circles) and 54 healthy controls (depicted by black squares). *P* values were calculated using two-tailed unpaired t-tests comparing the geometric mean of CD25^high^FOXP3^+^ Tregs between T1D patients and healthy controls (HC).

**Supplementary Fig. 3.**
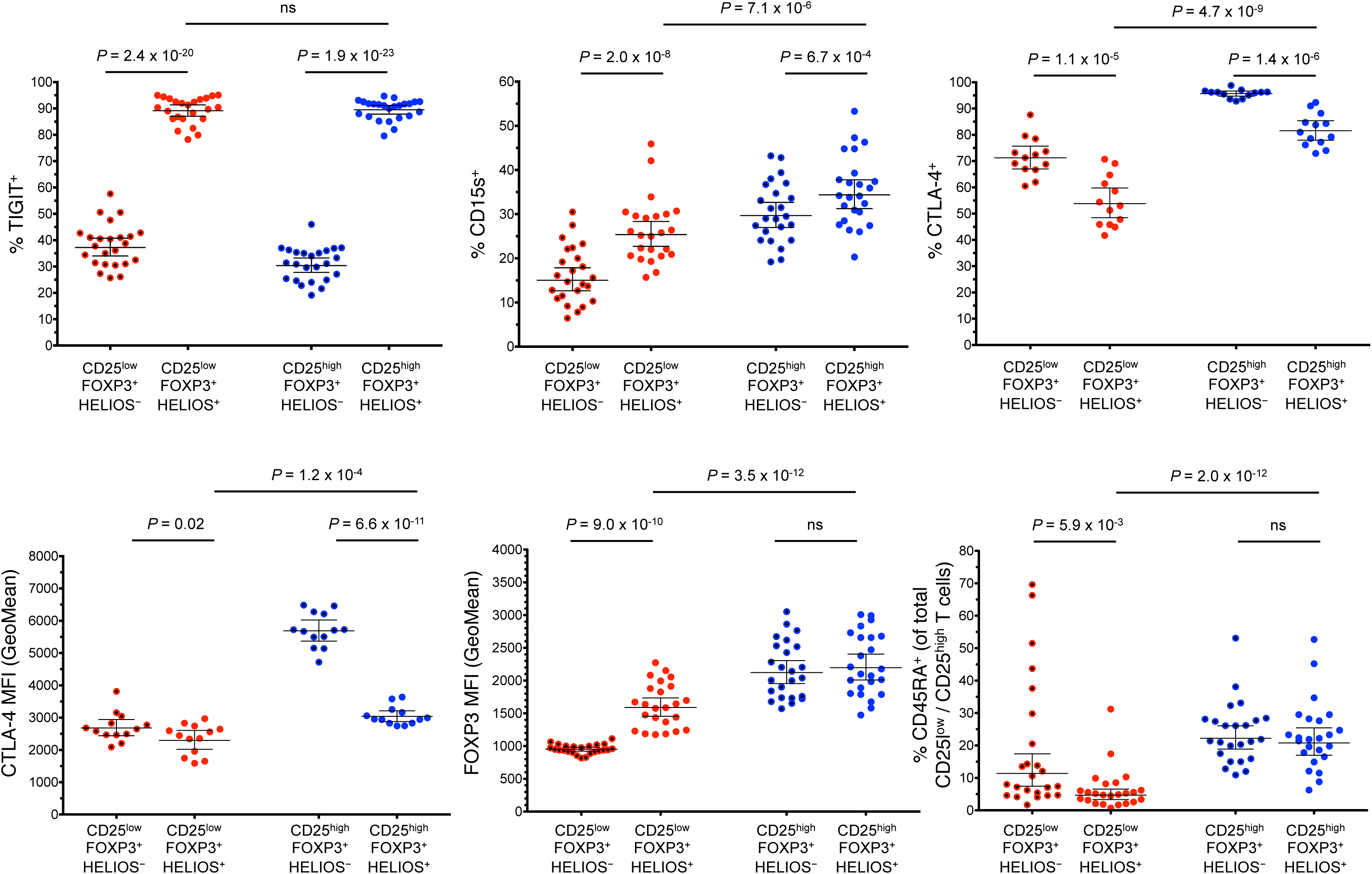
HELIOS expression defines distinct FOXP3^+^ subsets. Scatter plots depict the distribution (geometric mean +/- 95% CI) of TIGIT (n = 24), CD15s (n = 24), CD45RA (n = 24), CTLA-4 (both frequency and MFI of the positive fraction; n = 13) and FOXP3 MFI (n = 24) in the HELIOS^+^ and HELIOS^-^ fractions of the (i) CD25^low^FOXP3^+^ T cells (depicted in red) and (ii) conventional CD25^low^FOXP3^+^ Tregs (depicted in blue). *P* values were calculated using two-tailed paired t-tests.

**Supplementary Fig. 4.**
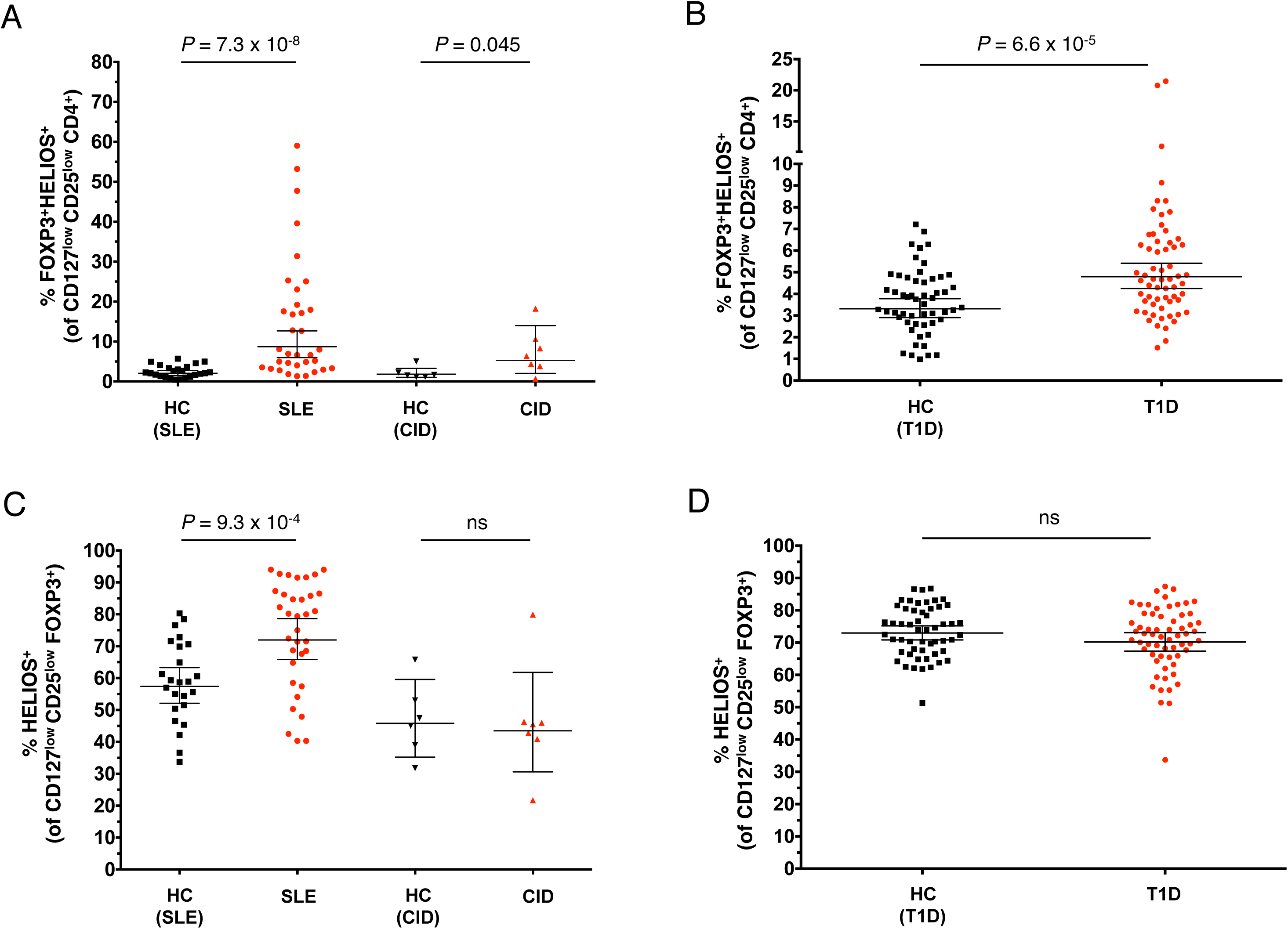
The frequency of HELIOS^+^ CD25^low^FOXP3^+^ cells is increased in patients with autoimmune disease. (**A, B**) Scatter plots depict the distribution (geometric mean +/- 95% CI) of HELIOS^+^FOXP3^+^ cells among CD127^low^CD25^low^ T cells in SLE patients (N = 32 patients vs 24 healthy donors) and combined immunodeficiency (CID) patients with active autoimmunity (N = 7 patients vs 6 healthy donors) (**A**); and in a cohort of T1D patients (N = 62; depicted by red circles) and healthy donors (N = 54; depicted by black squares) (**B**). (**C, D**) Scatter plots depict the distribution (geometric mean +/- 95% CI) of HELIOS^+^ cells within CD25^low^FOXP3^+^ T cells in the cohort of SLE and CID patients (**C**) and in the cohort of T1D patients (**D**). *P* values were calculated using two-tailed unpaired t-tests comparing the geometric mean of the assessed immune subsets between patients and the respective healthy control groups. .HC, healthy controls; T1D, type 1 diabetes patients; SLE, systemic lupus erythematosus patients; CID, combined immunodeficiency patients; ns = non-significant.

**Supplementary Fig. 5.**
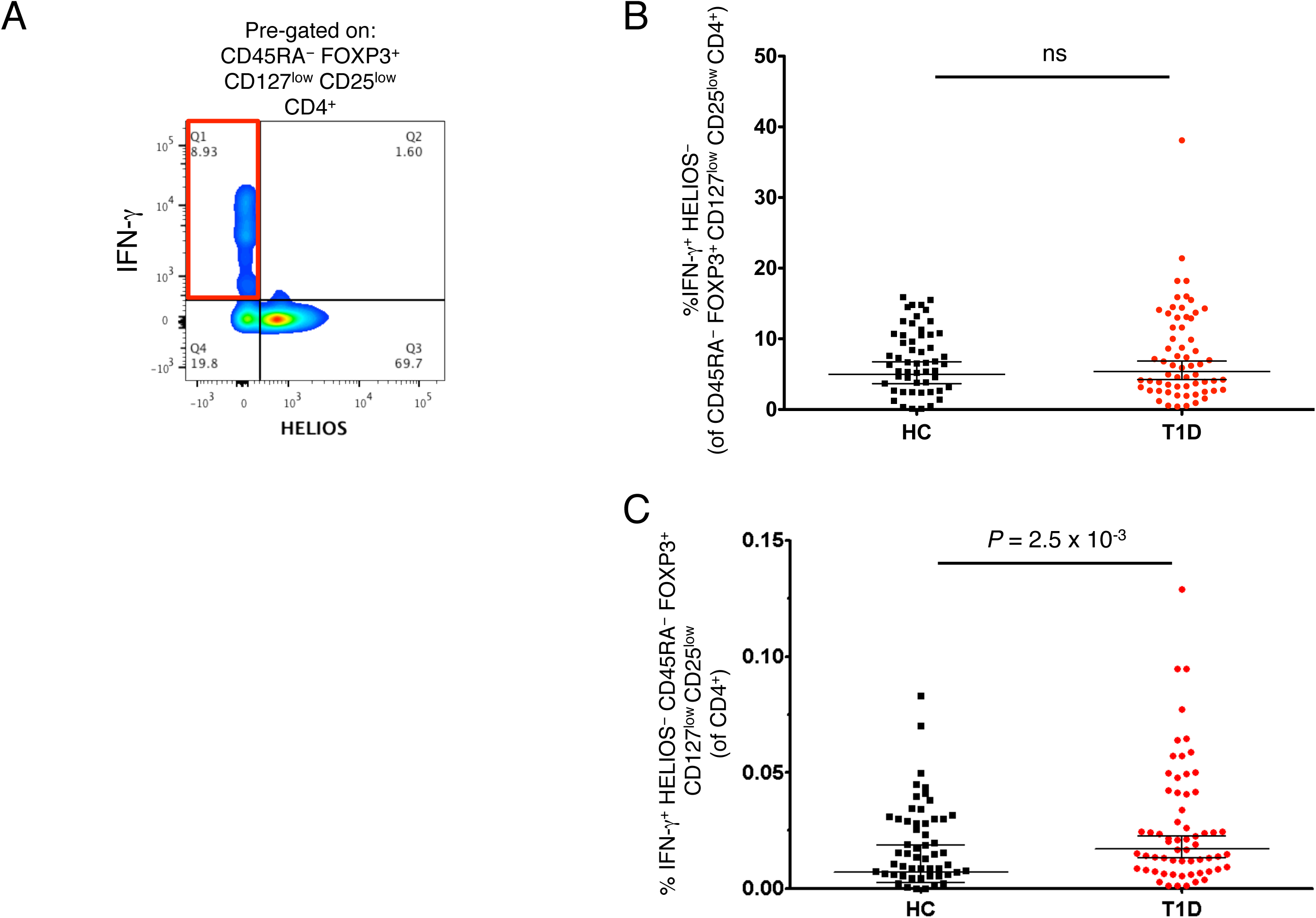
Production of IFN-γ from HELIOS^-^CD45RA^-^ CD127^low^CD25^low^FOXP3^+^ T cells is not altered in T1D patients. **(A)** Gating strategy illustrating the production of IFN-γ in the HELIOS^-^ and HELIOS^+^ CD45RA^-^ fractions of CD127^low^CD25^low^FOXP3^+^ cells. FACS gating plot is a representative example. **(B)** Plot depicts the distribution of the frequency (geometric mean +/- 95% CI) of IFN-γ+ HELIOS^-^ T cells in the CD45RA^-^ CD127^low^CD25^low^FOXP3^+^ population. Frequency of IFN-γ+ cells was compared between T1D patients (N = 62; depicted by red circles) and healthy donors (N = 54; depicted by black squares) following *in vitro* stimulation with phorbol-12-myristate-13-acetate (PMA) and ionomycin. (**C**) Plot depicts the distribution of the frequency (geometric mean +/- 95% CI) of IFN-γ+ HELIOS^-^ T cells in the CD45RA^-^ CD127^low^CD25^low^FOXP3^+^ population out of total CD4 T cells from the same donors as in (**B**). *P* values were calculated by linear regression of the log-transformed data, including batch as a covariate. HC, healthy controls; T1D, type 1 diabetic patients.

**Table 1.**
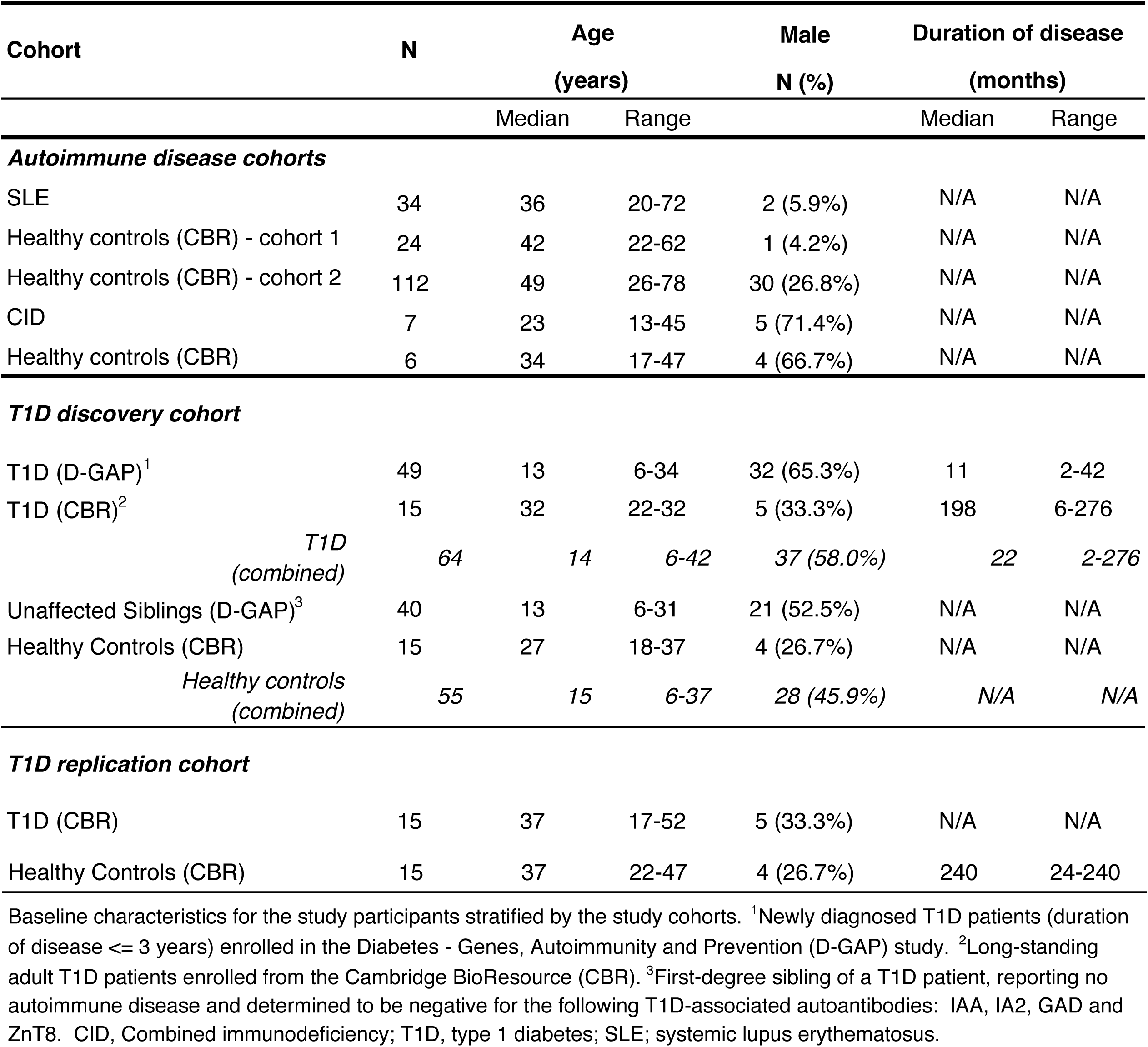
Baseline characteristics of study participants included in the association analyses

| Cohort | N | Age |  | Male | Duration of disease |  |
| --- | --- | --- | --- | --- | --- | --- |
|  |  | (years) |  | N (%) | (months) |  |
|  |  | Median | Range |  | Median | Range |
| ***Autoimmune disease cohorts*** |  |  |  |  |  |  |
| SLE | 34 | 36 | 20-72 | 2 (5.9%) | N/A | N/A |
| Healthy controls (CBR) - cohort 1 | 24 | 42 | 22-62 | 1 (4.2%) | N/A | N/A |
| Healthy controls (CBR) - cohort 2 | 112 | 49 | 26-78 | 30 (26.8%) | N/A | N/A |
| CID | 7 | 23 | 13-45 | 5 (71.4%) | N/A | N/A |
| Healthy controls (CBR) | 6 | 34 | 17-47 | 4 (66.7%) | N/A | N/A |
| ***T1D discovery cohort*** |  |  |  |  |  |  |
| T1D (D-GAP)<sup>1</sup> | 49 | 13 | 6-34 | 32 (65.3%) | 11 | 2-42 |
| T1D (CBR)<sup>2</sup> | 15 | 32 | 22-32 | 5 (33.3%) | 198 | 6-276 |
| *T1D*<br>*(combined)* | *64* | *14* | *6-42* | *37 (58.0%)* | *22* | *2-276* |
| Unaffected Siblings (D-GAP)<sup>3</sup> | 40 | 13 | 6-31 | 21 (52.5%) | N/A | N/A |
| Healthy Controls (CBR) | 15 | 27 | 18-37 | 4 (26.7%) | N/A | N/A |
| *Healthy controls*<br>*(combined)* | *55* | *15* | *6-37* | *28 (45.9%)* | *N/A* | *N/A* |
| ***T1D replication cohort*** |  |  |  |  |  |  |
| T1D (CBR) | 15 | 37 | 17-52 | 5 (33.3%) | N/A | N/A |
| Healthy Controls (CBR) | 15 | 37 | 22-47 | 4 (26.7%) | 240 | 24-240 |

Baseline characteristics for the study participants stratified by the study cohorts. <sup>1</sup>Newly diagnosed T1D patients (duration of disease ≤ 3 years) enrolled in the Diabetes - Genes, Autoimmunity and Prevention (D-GAP) study. <sup>2</sup>Long-standing adult T1D patients enrolled from the Cambridge BioResource (CBR). <sup>3</sup>First-degree sibling of a T1D patient, reporting no autoimmune disease and determined to be negative for the following T1D-associated autoantibodies: IAA, IA2, GAD and ZnT8. CID, Combined immunodeficiency; T1D, type 1 diabetes; SLE; systemic lupus erythematosus.

